# Sub-clustering, marker identification and multi-omics integration influenced by protoplasting in plant scRNA-seq data analysis

**DOI:** 10.1101/2024.12.14.628512

**Authors:** Jie Yao, Nianmin Shang, Hongyu Chen, Yurong Hu, Qian-Hao Zhu, Longjiang Fan, Chuyu Ye, Qinjie Chu

**Affiliations:** Institute of Crop Science, Zhejiang University, Hangzhou 310058, China; School of Medicine, Hangzhou City University, Hangzhou, China; CSIRO Agriculture and Food, Canberra, ACT 2601, Australia

**Author notes:** To whom correspondence should be addressed: Qinjie Chu.

**Keywords:** plant scRNA-seq, protoplasting effects, expression bias, protoplasting-induced genes

## Abstract

Single-cell RNA sequencing (scRNA-seq) technology is a powerful tool for exploring cell heterogeneity and lineage dynamics, revealing complex mechanisms driving tissue function and development. However, plant cell’s rigid cell walls require a protoplasing step, an enzymatic process that can introduce biases. The impact of protoplasting on plant scRNA-seq data remains poorly understood. In this study, we analyzed gene expression patterns from bulk RNA-seq of various plant tissues before and after protoplasting. Cell-type specific protoplasting effects were discovered in different plant tissues. Protoplasting-related biases were found to distort clustering, marker gene identification, and multi-omics integration. By calculating protoplasted scores based on gene expression, we identified and mitigated protoplasting effects, enabling the accurate clustering and annotation of tobacco BY-2 cells and identification of cell-cycle-related genes. Our findings underscore the importance of assessing and correcting for protoplasting biases in plant scRNA-seq data analysis, offering new insights for more accurate data interpretation and biological discovery.

## Introduction

Single-cell RNA sequencing (scRNA-seq) has revolutionized our understanding of cellular heterogeneity and lineage dynamics, providing deep insights into the functions and developmental processes of diverse tissues. This cutting-edge technology has been extensively used to explore the complex molecular landscapes of various plant species and tissues ^1, 2^, including *Arabidopsis thaliana* ^3–5^, *Oryza sativa* ^6^, *Zea mays* ^7^, *Solanum lycopersicum* ^8, 9^, *Populous* ^10^, *Fragaria vesca* ^11^, *Nicotiana* ^12^, and so on. These studies have significantly enhanced our knowledge of cellular diversity, developmental processes, and response mechanisms of these plant species.

Despite significant advancements in scRNA-seq studies across various plant species, the impact of enzymatic digestion induced by protoplasting on data analysis remains under explored, limitted the accurate data interpretation. Additionaly, the varying conditions and durations required for protoplasting across plant species and tissue types present further challenges during pre-processing stage ^13^ and subsequent data analysis ^14^.

To address this knowledge gap, we analyzed public bulk RNA-seq datasets of various plant tissues before and after protoplasting to identify <u>p</u>roto<u>p</u>lasting-related <u>d</u>ifferentially <u>e</u>xpressed <u>g</u>enes (ppDEGs). We quantified the effects of enzymatic digestion in single cells using <u>p</u>rotoplasted <u>s</u>core (ppScore) values, which effectively capture the protoplasting effects in plant scRNA-seq data. Our analysis revealed cell-type-specific protoplasting effects, which could influence sub-clusting and marker gene identification. Applying ppScore values to scRNA-seq data from tobacco BY-2 cells, we successfully annotated cell-cycle-related cell types. By highlighting enzymatic hydrolysis biases induced by protoplasting, this study provides a deeper understanding of the challenges and opportunities in decoding the molecular landscape of plant tissues at the single-cell level, advancing our knowledge for plant development, physiology, and evolutionary biology.

## Results

### Gene expression significantly changed while protoplasting

To investigate gene expression changes caused by protoplasting, bulk RNA-seq datasets (**Table S1**) generated from plant tissues and corresponding protoplasts were collected to identify <u>p</u>roto<u>p</u>lasting related <u>d</u>ifferentially <u>e</u>xpressed <u>g</u>enes (ppDEGs). Briefly, a total of 3 435, 5 878, 5 850 and 9 725 ppDEGs were identified from the root and leaf of *A. thaliana*, *Z. mays* leaf and *O. sativa* seedling RNA-seq samples (**Table S1, S2**), respectively. Expression correlation of datasets from scRNA-seq (considered as pseudo bulk RNA-seq), un-protoplasted and protoplasted RNA-seq showed that samples by scRNA-seq and protoplasted RNA-seq had a more similar expression pattern than by un-protoplasted samples (**Fig. S1A-C**, R^2^ 0.72 *vs* R^2^ 0.22, R^2^ 0.62 *vs* R^2^ 0.17, R^2^ 0.56 *vs* R^2^ 0.33 in *A. thaliana*, *O. sativa* and *Z. mays* respectively). These results revealed that gene expression values changed variously in the process of protoplasting and such diversity was also maintained in the scRNA-seq dataset because of the necessity of cell dissociation.

### Cell-type specific protoplasting effects uncovered by scRNA atlases

To elucidate the specific impacts of protoplasting across different cell types in diverse plant species, we generated integrated single-cell transcriptome atlases for root (**Fig. 1**), leaf (**Fig. S2A-C**), and flower tissues (**Fig. S2D-F**) from *A. thaliana*. The *A. thaliana* root atlas (**Fig. 1A**), encompassing over 160k cells, consisted of 14 cell types organized within continuous branches corresponding to five tissues ^15^. Protoplasting effects were quantified as ppScore (<u>p</u>roto<u>p</u>lasted <u>score</u>) values, which were calculated by the *addModuleScore* function in Seurat ^16^ based on the expression of ppDEGs. For *A. thaliana* root cells, compared with other cell types, significantly higher (Wilcoxon rank sum test, *p*-value < 0.001) ppScore values were observed in the root cap, including lateral root cap (cluster 0, 10, 12) and columella cells (cluster 3, 24, 25, 28) (**Fig. 1B-C**). This suggested that root cap was particularly vulnerable to protoplasting stimulation. We also performed the same analysis in root (**Fig. 2**), leaf (**Fig. S3A-C**) and inflorescence tissues (**Fig. S3D-F**) of *O. sativa*. The scRNA atlas of *O. sativa* root, comprising over 86k cells, was annotated into 15 cell types and classified into four cell lineages (**Fig. 2A**). Results showed that vascular cylinder cells (cluster 4, 5, 6, 15, 20, 23, 24, 25, 30) had higher ppScore values (Wilcoxon rank sum test, *p*-value < 0.001, **Fig. 2B-C**) compared with other cell types, which were considered as cell types that were susceptible to protoplasting.

**Fig. 1.**
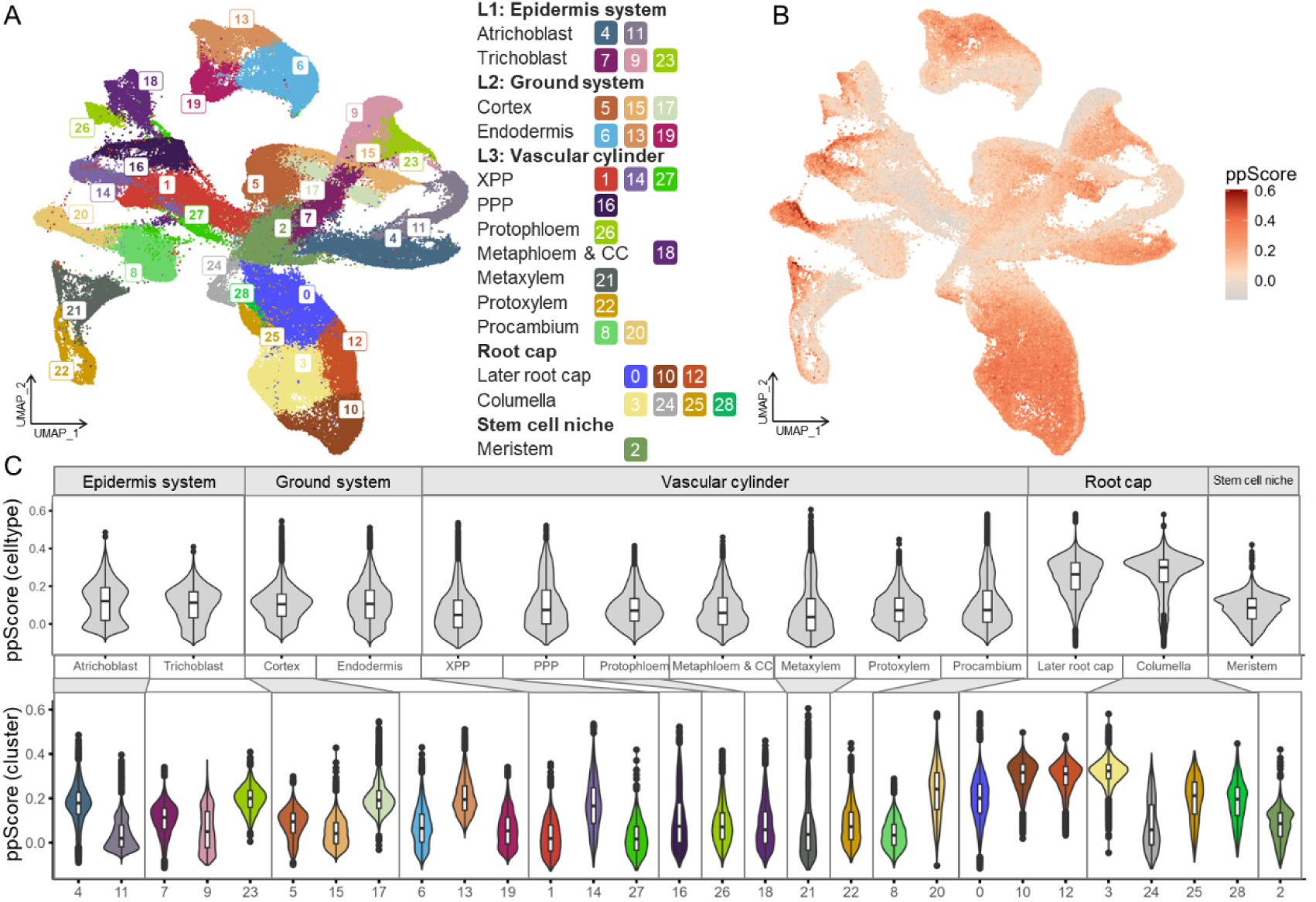
Construction of *A. thaliana* root scRNA atlas and quantification of protoplasting effects by ppScore values in different cell types and cell clusters. **A:** Integration of scRNA-seq datasets from *A. thaliana* roots (PRJNA509920, PRJNA640389), which comprised over 160k cells across 14 cell types. **B:** Visualization of ppScore values with darker colors indicating higher protoplasting impacts. **C:** Variations of ppScore values across different cell types (upper) and cell clusters (bottom).

**Fig. 2.**
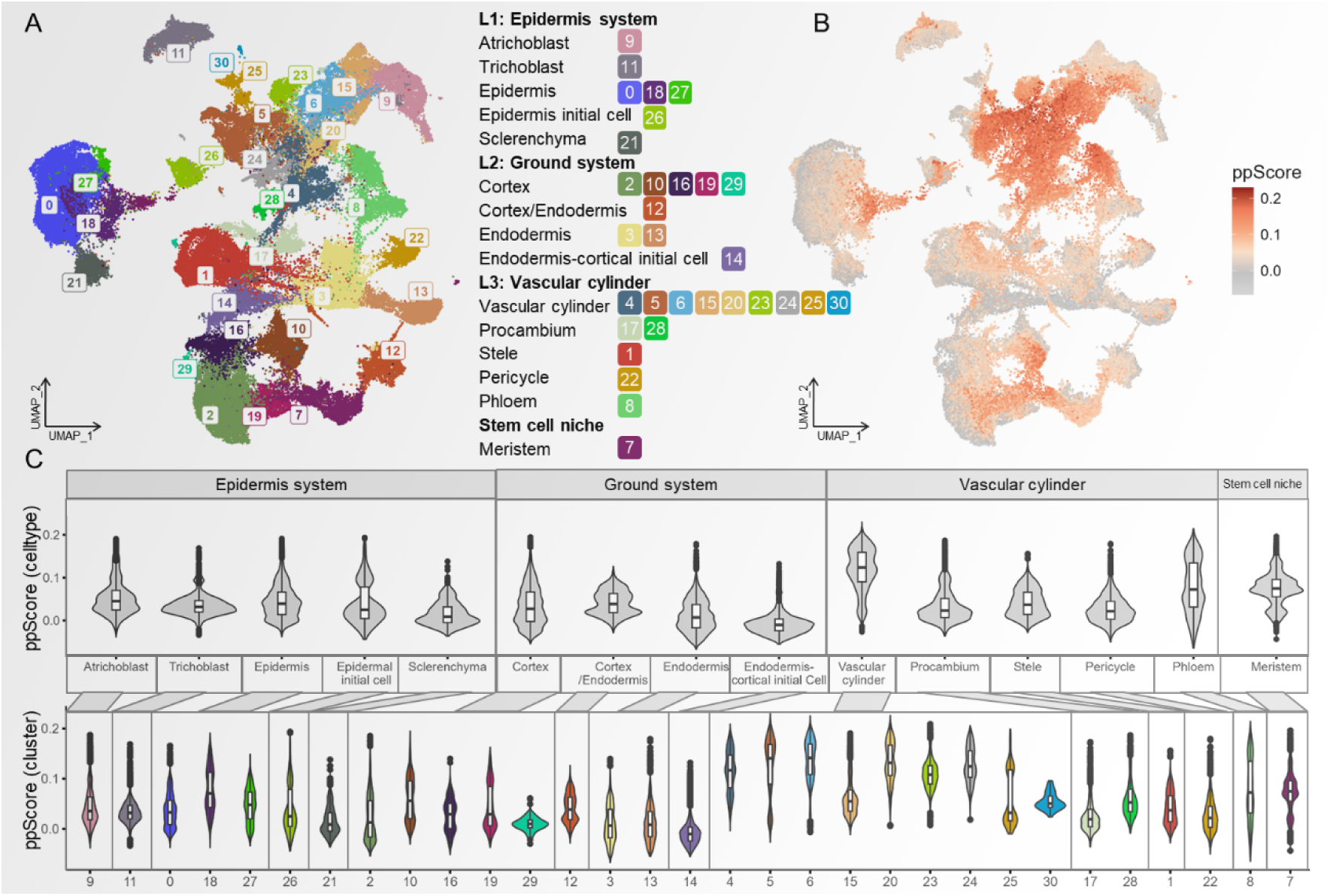
Construction of *O. sativa* root scRNA atlas and quantification of protoplasting effects by ppScore values in different cell types and cell clusters. **A:** Integration of scRNA-seq datasets from *O. sativa* roots (PRJNA706435, CRA004082), which comprised over 86k cells across 15 cell types. **B:** Visualization of ppScore values with darker colors indicating higher protoplasting impacts. **C:** Variations of ppScore values across different cell types (upper) and cell clusters (bottom).

Furthermore, single-cell transcription comprehensive atlases of *A. thaliana* leaf (**Fig. S2A-C**) and floral receptacles (**Fig. S2D-F**), *O. sativa* leaf (**Fig. S3A-C**) and flower (**Fig. S3D-F**), all demonstrated similar results that cells with strong growing ability (such as young vascular cells in *O. sativa* leaf) and differentiation ability (such as meristem cells in *A. thaliana* leaf and *O. sativa* flower) were affected greatly by protoplasting, which can provide clues for further investigation.

**Cells response to protoplasting were clustered as sub-type within the same cell type** Furthermore, different cell clusters within the same cell type were observed with significant variations of ppScore values (**Fig. 1C, 2C**). For example, in the epidermis system of *A. thaliana root* scRNA atlas, clusters 4 and 23 demonstrated higher (Wilcoxon rank sum test, *p*-value < 0.001) ppScore values than other clusters in atrichoblast and trichoblast cells, respectively (**Fig. 1C**). Additionally, cluster 17 in cortex and cluster 13 in endodermis cells in the ground tissue, cluster 3 of columella cells experienced more obvious protoplasting stimulation with higher ppScore values than other cell clusters (all *p*-value < 0.001 by Wilcoxon rank sum test). Similarly, from *O. sativa* root scRNA atlas, cluster 18 of epidermis cells and cluster 10 of cortex cells were of highest ppScore values than the other clusters in the same cell types (**Fig. 2C**, *p*-value < 0.001 by Wilcoxon rank sum test). Interestingly, although meristem cells had only one cluster (cluster 7), the bimodal distribution of ppScore values was clearly distinguished by the violin plot (**Fig. 2C**). The results showed that protoplasting influenced sub-type clustering within the same cell types by higher expression of ppDEGs with higher values of ppScore.

In order to investigate the clustering differences of some cell types before and after removing protoplasting effects, three methods were employed including, 1) Method One (M1): removing protoplasting biases by *ScaleData* function in R package Seurat; 2) Method Two (M2): directly removing cells with high ppScore values, the threshold value of which is determined by users, usually could be set to 0 or 0.1; 3) Method Three (M3): directly remove ppDEGs from raw expression matrixes. Results showed that in trichoblast cells of *A. thaliana* root, with the same analysis parameters, cells with high ppScore values were mainly clustered in cluster 0 and 2 in raw data, but these cells were separated into three clusters after removing protoplasting effects by M1 and M3 (**Fig. 3**). The results indicated the effectiveness of removing protoplasting effects, and cells seriously affected by protoplasting were clustered relatively dispersedly after removing protoplasting effects. However, clustering results of different cell types from different plant species and tissues was quite not the same after removing protoplasting effects (**Fig. S4**), with some demonstrated the effectiveness of removing methods while others show no significant changes. Of course, removing high ppScore cells (M2) was the most direct and convenient method for dealing with protoplasting effects, but might face a risk of losing important biological cells.

**Fig. 3.**
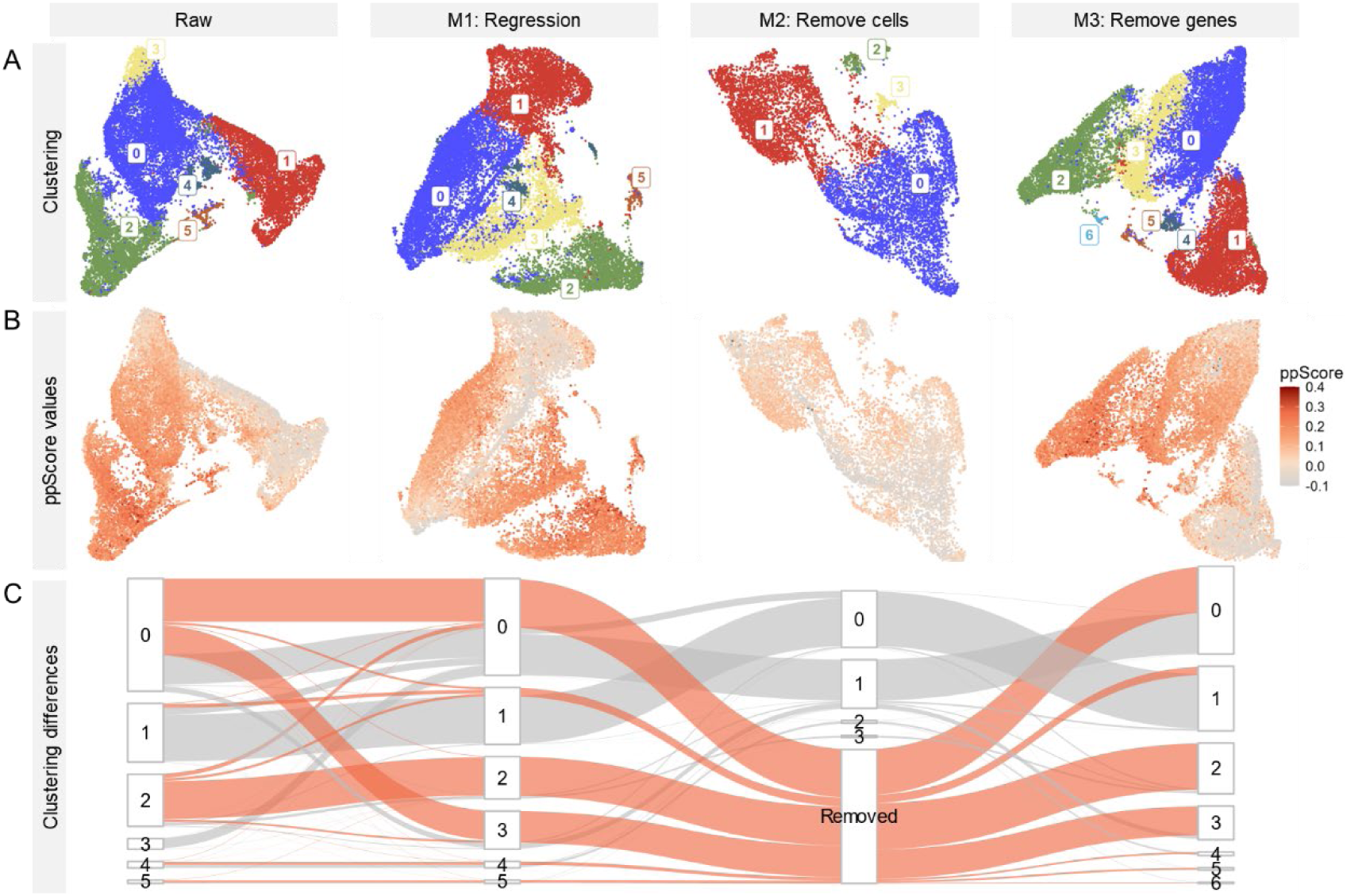
Clustering differences of *A. thaliana* trichoblast cells between raw and modified datasets (i.e. removing protoplasting effects by three methods) with same parameters (res = 0.1). **A:** UMAP visualization of clustering results for raw and three modified datasets. Three methods to remove protoplasting effects were detailly described in main text. **B:** Values of ppScore in different datasets, while darker color of points represents higher ppScore value, indicating higher protoplasting impacts. **C:** Sankey plot showed the changes of cell clustering among different datasets. Orange lines represent cells with high ppScore values (in this case, ppScore > 0.1) that were removed by Method Two (M2).

### Marker gene identification differences caused by protoplasting

Intersection between ppDEGs and marker genes of each cell cluster were investigated to discover the identification differences of marker genes caused by protoplasting. Results showed that the proportion of ppDEGs in all identified marker genes (logFC >= 0.25 and adjusted p-value ⩽ 0.05, **Fig. 4A, 4E, S4**) was consist with the distribution of ppScore values(**Fig. 1C, 2C, S2C, S2F, S3C, S3F**) of each cell cluster, that’s to say, cell clusters with higher ppScore values usually had higher proportion of ppDEGs. The values of ppDEG proportion were related to the number of ppDEGs (**Table S1**). Because there are nearly 6000 ppDEGs identified in *O. sativa* but only about 1000 ppDEGs in *A. thaliana* root, the proportion of ppDEGs in *O. sativa* (**Fig. 4E**) was higher than that of *A. thaliana* (**Fig. 4A**). Besides, we also investigated the intersection of protoplasting-related genes identified as marker genes in sub-type analysis, before and after removing protoplasting effects by three methods as described in the previous section (**Fig. 4B-D, 4F-H**). Results showed that, to some extent, removing protoplasting effects could reduce the ppDEG proportion in some cell types (**Fig. 4B-D, H**) and maintain the majority of the intersection. However, in different cell types, the phenomena may be different, which need a concrete analysis of specific situations.

**Fig. 4.**
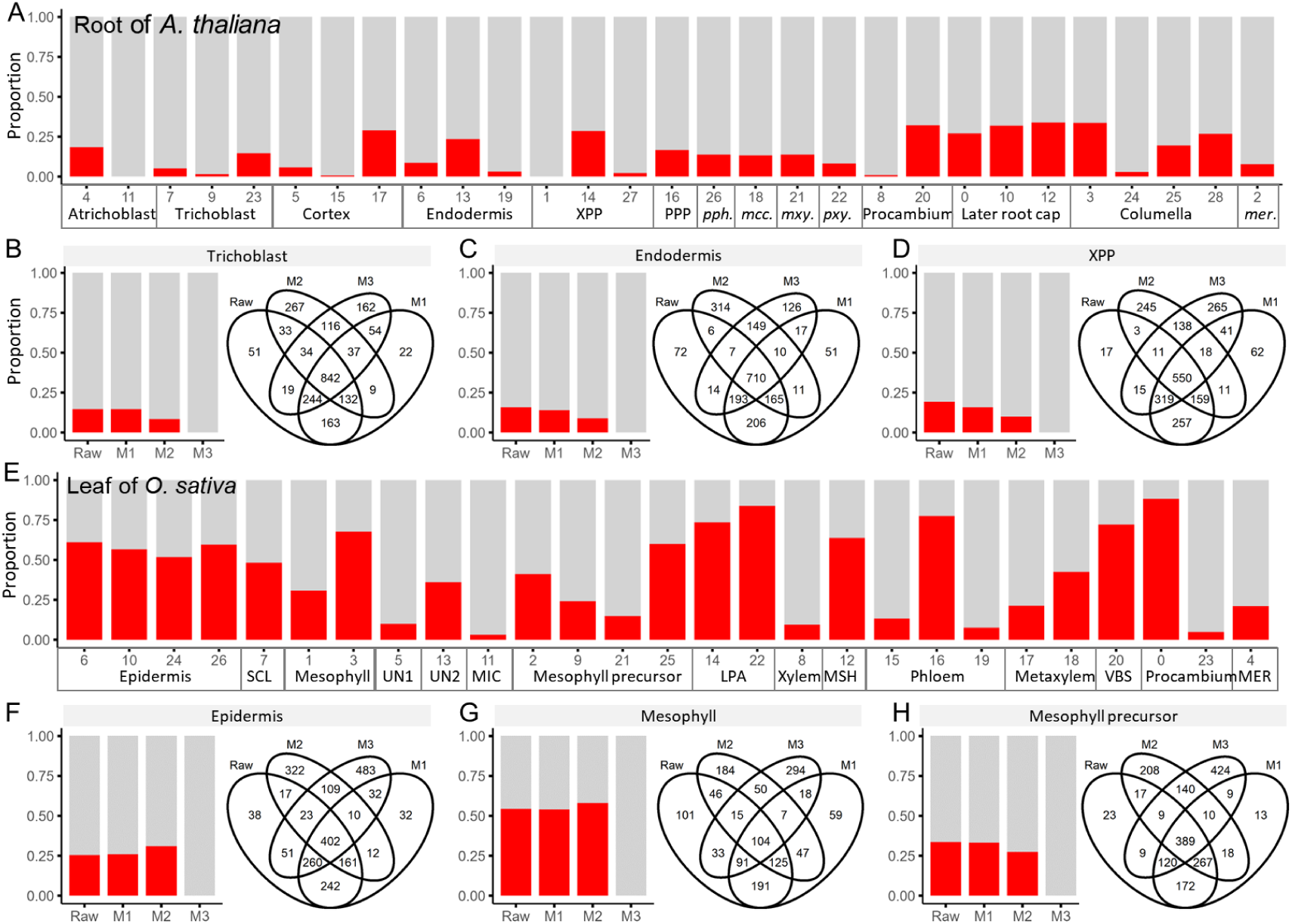
Proportion of ppDEGs in marker genes of each cell cluster in *A. thaliana* root (A-D) and *O. sativa* leaf (E-H). **A:** Proportion of ppDEGs (red bars) in each cell clusters of *A. thaliana* root. **B-D:** Intersection and ppDEGs proportion in raw and modified datasets (i.e. removing protoplasting effects by three methods) of trichoblast (**B**), endodermis (**C**) and XPP (**D**) cells, respectively. M1, M2, M3 represent to remove protoplasting effects by *ScaleData* function in Seurat, removing high ppScore cells and removing ppDEGs, respectively, which were detailly described in main text. **E:** Proportion of ppDEGs (red bars) in each cell clusters of *O. sativa* leaf. **F-G:** Intersection and ppDEGs proportion in raw and modified datasets of epidermis (**F**), mesophyll (**G**) and mesophyll precursor (**H**) cells, respectively.

### Integration of multi-omics data was influenced by protoplasting effects

Taking snRNA-seq and spatial transcriptome (ST) datasets, we aimed to illustrate the influences of protoplasting effects while integrating plant multi-omics data. Integrated analyses of scRNA-seq (PRJNA509920) and snRNA-seq (PRJNA649267) datasets from *A. thaliana* roots ^17, 18^, revealed ten cell types (**Fig. 5A**). Most cell types comprise commensurate cells derived from both scRNA-seq and snRNA-seq (**Fig. 5B**). However, a distinct cluster comprising cells extensively derived from scRNA-seq displayed no obvious known cell type marker genes but with prominently elevated ppScore values, posing challenges for annotation, and was thus labeled as “unknown” (**Fig. 5B**). Additionally, atrichoblast cells also exhibited a substantial elevation in ppScore values. Marker genes for these two highly impacted cell types (atrichoblast and unknown cells) were identified and GO (Gene Ontology) enrichment results of these genes revealed a significant enrichment of stress response-related pathways, such as cellular response to heat, response to decreased oxygen levels, response to wounding, and toxin metabolic processes (**Fig. 5C**).

**Fig. 5.**
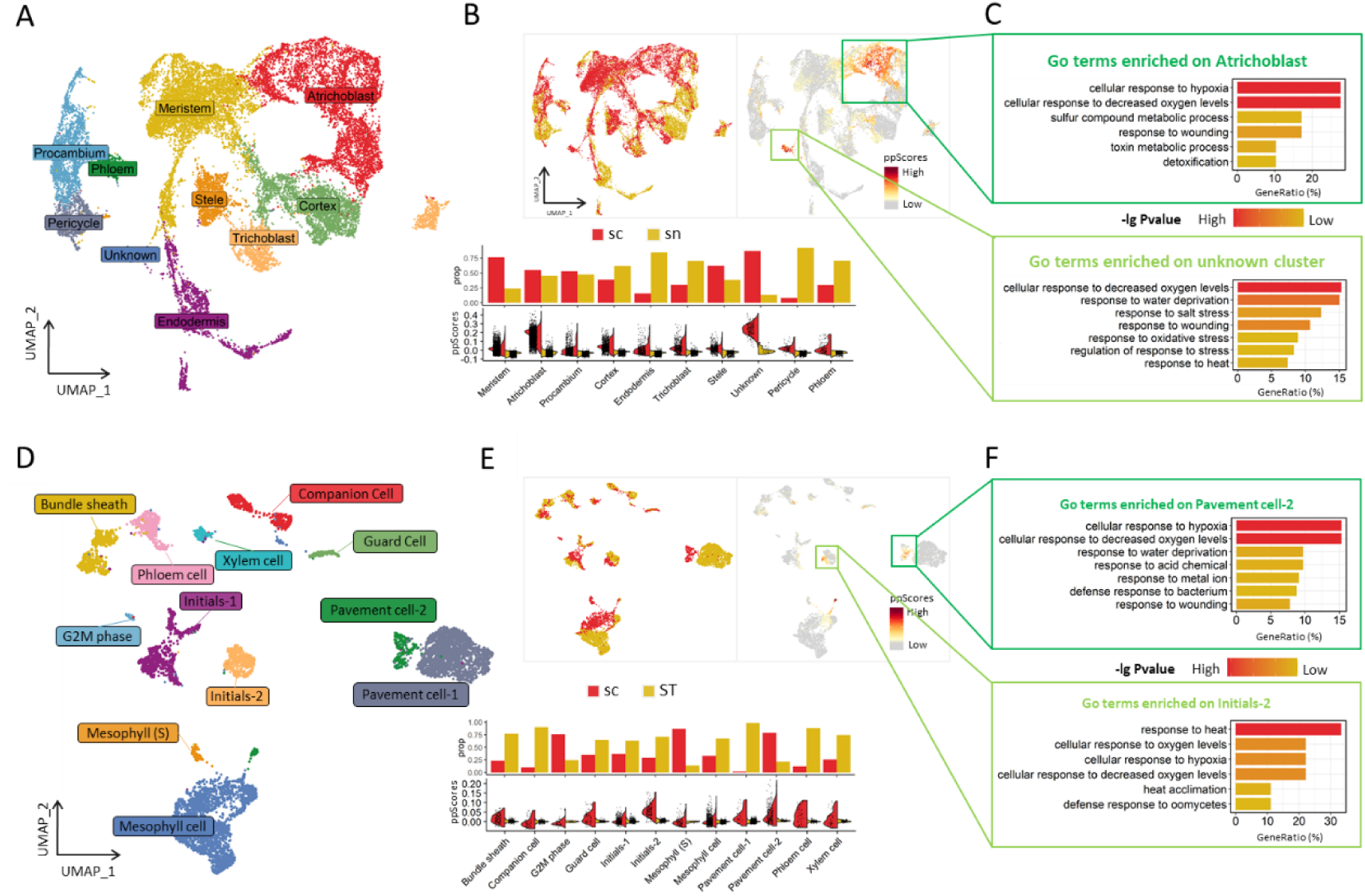
The existence of protoplasting effects in plant scRNA-seq data influenced the integration of single-cell multi-omics data. **A:** Integration of scRNA-seq and snRNA-seq datasets from *A. thaliana* roots, with ten different cell types identified and visualized by UMAP. Each dot denoted a single cell. **B:** The top left panel illustrated the UMAP result after the integration of scRNA-seq and snRNA-seq data, and the top right panel showed the expression patterns of ppDEGs. The bottom panel depicted the cell proportions from scRNA-seq or snRNA-seq data in each cluster and the overall ppScore values for each cluster, respectively. **C:** Representative GO (Gene Ontology) terms enriched in marker genes of cell types. **D:** Integration of scRNA-seq and ST datasets of *A. thaliana* leaves, with twelve cell types were identified and visualized by UMAP. **E-F** were analogous to **B-C**.

Furthermore, the integration dataset of scRNA-seq (ETAB-10016MT) and ST (CNP0002618) datasets from *A. thaliana* leaves ^19, 20^ was annotated with twelve distinct cell types (**Fig. 5D**). Similarly, in this dataset, we observed significantly higher ppScore values indicating higher protoplasting effects in cells originating from scRNA-seq compared to those from ST (**Fig. 5E**). The marker genes from the top two protoplasting-affected cell types (initials-2 and pavement cell-2) were enriched in pathways associated with heat response, decreased oxygen levels, water deprivation, wounding response, acid chemical response, metal ion response, and so on (**Fig. 5F**). Our results demonstrated the existence of protoplasting effects in scRNA-seq data may impact cell clustering while integrating with snRNA-seq or ST data, which should to be paid attention to.

### Overcoming protoplasting-induced effects for improved analysis of scRNA-seq data from tobacco BY-2 cells

For confirmation of the effectiveness and necessity for analyzing protoplasting-induced effects on plant scRNA-seq data, the tobacco BY-2 cell line was selected for scRNA sequencing because of its simple cellular composition and cell homogeneity. Protoplasts were isolated from three-day-old BY-2 suspension cells and a total of over 12,000 cells from two biological replicates were collected (**Fig. 6A**). The two BY-2 cell line samples exhibited good reproducibility (Spearman’s correlation = 0.90, **Fig. S6A**). Bulk RNA-seq from protoplasted and untreated BY-2 cells were also performed (three replicates for each condition), and 4,350 genes were classified as ppDEGs (|log2FC| ≥ 2, *p*-value ≤ 0.05). We observed that the pseudo-bulk scRNA-seq results of BY-2 cells (protoplasted cells) showed a low correlation with the unprotoplasted bulk mRNA expression (Spearman’s correlation = 0.69, **Fig. S6B**), which was consistent with the results shown by **Fig. S1**. Finally, a pool of 11,775 high-quality cells (7,443 and 4,332 from two replicates, respectively) were selected, which expressed a total of 61,657 genes, for further analysis.

**Fig. 6.**
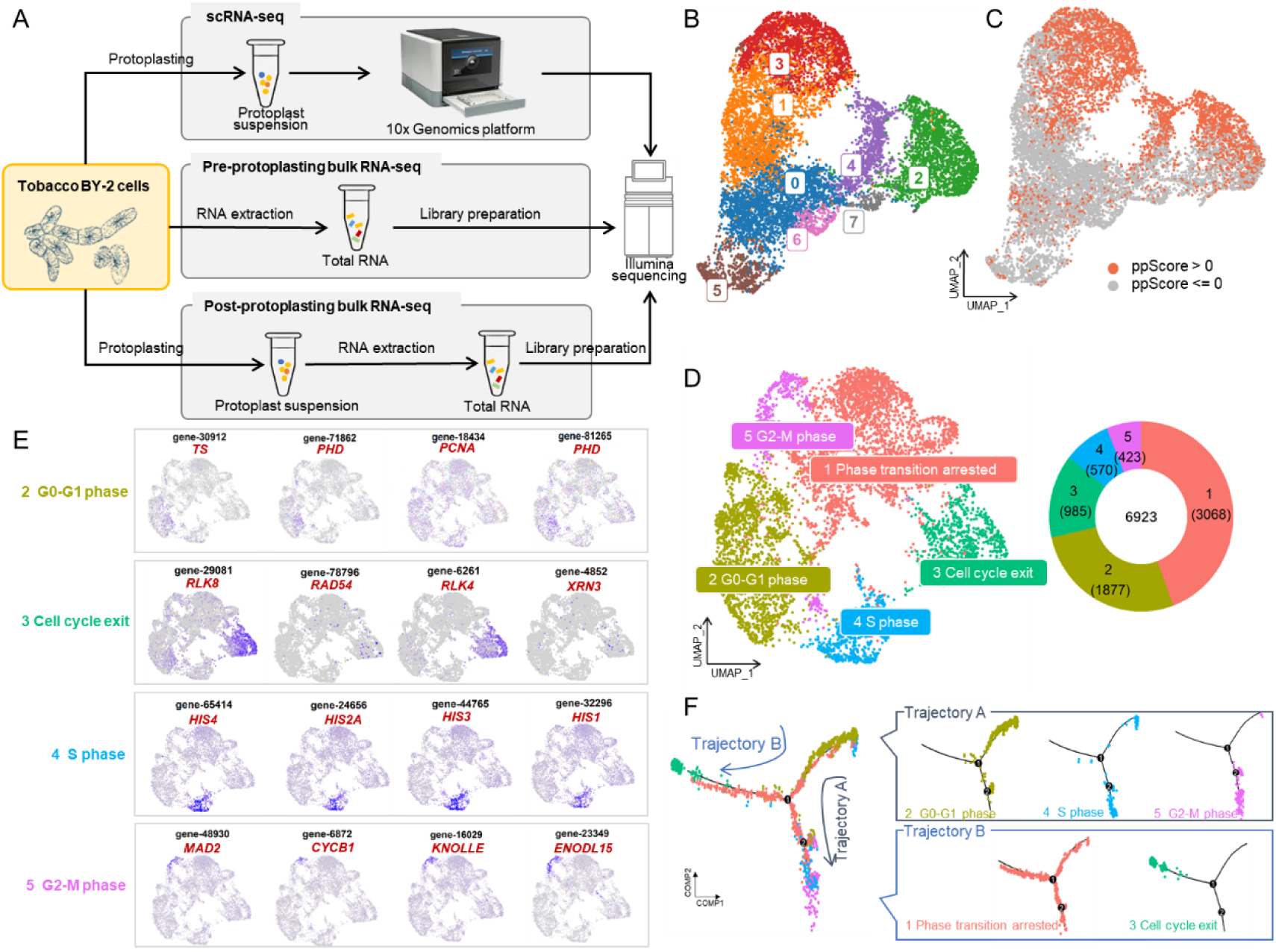
An example of applying ppScore values for analysis of scRNA-seq data from tobacco BY-2 cells. **A:** A schematic showed the experimental design of tobacco BY-2 cells. Details were described in *Materials and Methods.* **B:** UMAP plot containing 11,775 BY-2 cells were grouped into eight clusters. Distinct clusters were indicated by different colors and dots represented individual cells. **C:** Distribution of cells with high (red dots, 4 852 cells, ppScore > 0) and low (blue dots, 6 923 cells, ppScore <= 0) ppScore values, which indicating cells were highly and lowly affected by protoplasting, respectively. **D:** Visualization of the five cell types clustered by using 6 923 BY-2 cells after removing cells with high ppScore values. The right panel showed the cell number of each cluster. **E:** Expression pattern of selected cell cycle phase-specific genes in G0-G1 phase, S phase, G2-M phase, and cell cycle exit. **F:** Simulation of the successive differentiation trajectory of BY-2 cells by pseudo-time. The right panel showed the relationship between clusters (1-5) and trajectories (indicated by arrows).

Unsupervised analyses grouped the BY-2 cells into 8 distinct clusters (**Fig. 6B**). By calculating ppScore values, approximately 41.2% (4,852 out of 11,775) of all cells showed a high degree of protoplasting impact (ppScore > 0), predominantly located in the upper region of the UMAP plot where clusters 1, 4, and 5 were situated (**Fig. 6C**). Out of 4,087 identified marker genes of each cell cluster, 714 (17.5%) genes overlapped with ppDEGs. By excluding cells with high ppScore values and reanalyzing the remaining 6,923 cells, five new clusters were identified (**Fig. 6D**, detailed described in *Methods*). Each cluster was assigned to a specific phase or characteristic of the cell cycle based on gene expression profiles and known cell cycle-related genes (**Fig. 6E**). Moreover, pseudo-time trajectory analysis was performed by using cluster marker genes to explore the cell cycle progression of tobacco BY-2 cells (**Fig. 6F**). Results showed that the trajectory began in cluster 2, representing the G0-G1 phase, and branched into two trajectories indicating different cell fates. Trajectory A represented the progression from G0-G1 phase (cluster 2) to S phase (cluster 4) and G2-M phase (cluster 5). Trajectory B included cluster 3 and part of cluster 1, which might represent an exceptional case of cell cycle phase transition, such as cells arrested or exited due to DNA replication damage or environmental cues. The trajectory analysis not only enabled the reconstruction of cell fate differentiation but also confirmed the accuracy of our cell identity assignment in the analysis.

This example of single cell transcriptomes from tobacco BY-2 cells displayed a good story of cell division progression and cell cycle exit after evaluating of protoplasting effect and removing of cell strongly affected by protoplasting, which indicated the importance of ppScore values for protoplasting effect evaluation.

## Discussion

In this study, we systematically investigated the impact of enzymatic digestion biases caused by protoplasting on plant scRNA-seq data analysis, with the help of bulk RNA-seq, snRNA-seq and ST data. The impact of ppDEGs on plant scRNA-seq analysis has been largely overlooked in previous studies. Among 53 published articles involving plant scRNA-seq technology, covering 11 species and over 15 tissues, only 12 studies (22.6%) explicitly addressed the potential enzymatic digestion biases introduced by protoplasting (**Table S3**). A better way to obtain the accurate set of DEGs influenced by protoplasting in different species or tissues is to conduct bulk RNA-seq before and after protoplasting, which is highly recommended by us. By calculating ppScore (<u>p</u>roto<u>p</u>lasted <u>score</u>) values which indicating the degree of protoplasting effects, we successfully applied it to different datasets from different plants and tissues (**Fig. 1, 2, S2, S3**). The influences of protoplasting may be different among different datasets, for example, the cell cluster with high ppScore values could not be properly annotated in **Fig. 5B**, but conversely could be assigned as initial and pavement cells in **Fig. 5E**.

Cell-type specific protoplasting effects were revealed in this study. Detailly, root cap cells in *A. thaliana* roots (**Fig. 1C**), young vascular cells or meristem cells in other investigated plants were found significantly affected by protoplasting (**Fig. 2, S2, S3**). For *A. thaliana* root cap cells, which are located at the forefront of the root system and possess a less dense cellular structure ^21^, because of their location and structure, they may respond more quickly, easily and greatly to protoplasting. *O. sativa* vascular cells shown in **Fig. 2**, which were surrounded by cells with differentiation ability such as procambium (cluster 17, 28) and epidermis initial cells (cluster 26), were relatively young or newly formed cells that might be sensitive to protoplasting. Vascular cells in

*A. thaliana* leaves and *O. sativa* leaves had a similar tendency (**Fig. S2C, S3C**). Therefore, considering and accounting for the influences caused by cell wall digestion is crucial when interpreting the biological significance of these particular cell types. Besides, with the accumulation of more plant scRNA-seq datasets and the construction of complete standard expression atlases of plants, cell-type specific protoplasting effects could be investigated in more plant species and tissues.

Interestingly, our study discovered that protoplasting effects displayed discernible variations among distinct cell clusters within each cell type, highlighting their profound influences on the characterization of cell subpopulations. Cell subpopulation analysis is common in single-cell RNA sequencing studies. However, results of sub-type could be influenced by protoplasting effects. For example, newly identified subpopulations may not solely represent biological differences but could instead result from biases introduced by protoplasting (**Fig. 1C, 2C**). Caution is necessary when defining subpopulations to avoid misclassifying clusters heavily impacted by protoplasting as distinct biological subtypes. Addtionaly, it is important to recognize that ppDEGs can affect not only clustering but also the identification of marker genes for specific cell clusters, potentially altering downstream biological interpretations of cell types.

In addition, enzymatic digestion leads to diverse expression patterns of differentially expressed genes, influenced by variations in tissue maturity or protoplasting conditions. For example, in maize ear data ^22^, only 732 up-regulated DEGs (log2FC≥2, p_value≤0.05, much less than DEGs displayed in **Table S1**) were identified, as only 45 minutes of protoplasting was required for the tender ear tissue becoming intact protoplasts. In contrast, another maize leaf dataset ^23^ needed 210 minutes of protoplasting and yielded 2,765 up-regulated DEGs using the same criteria. On the other hand, the differences in sequencing platforms or depth, annotated genes of the reference genomes and other unnoticed factors all might influence the identified numbers of ppDEGs.

Furthermore, an essential consideration is the complementarity of multi-omics sequencing technologies, which could reduce information loss due to technical biases. While scRNA-seq is widely used, enzymatic protoplasting may introduce transcriptional biases in plant cells. Technologies of snRNA-seq and spatial transcriptomics offer alternatives to address this issue, with the former detecting nuclear-localized mRNAs and the latter preserving spatial information. However, snRNA-seq may miss certain genes or transcripts as it only detects nuclear-localized mRNAs ^24^. For spatial transcriptomics, its sequencing depth may be much lower than scRNA-seq data ^25^. Therefore, if possible, we suggest pursuing multi-omics sequencing at the single-cell level to achieve comprehensive information complementarity.

Overall, our study provides new insights into cellular heterogeneity and lineage dynamics in plant tissues at the single-cell level, highlightingpreviously unexplored enzymatic digestion biases in plant scRNA-seq analysis.

## Materials and Methods

### Identification of protoplasting related DEGs (ppDEGs)

Bulk RNA-seq reads were aligned to reference genomes by STAR version 2.7.9 ^26^. The reference genomes of *A. thaliana* (Tair10) and *Z. mays* (Maize V4) were downloaded from SOL genomics database (https://solgenomics.net/) and maizeGDB database (https://maizegdb.org/), respectively. Gene expression values were calculated by the number of uniquely mapped reads using featureCounts version 2.0.2 ^27^, and DEseq2 version 1.28.1 ^28^ was used to identify differentially expressed genes (DEGs). <u>P</u>roto<u>p</u>lasting related <u>d</u>ifferentially <u>e</u>xpressed <u>g</u>enes (ppDEGs) were identified by the criteria of |log2FC| ≥ 2, *p*-value ≤ 0.05 (FC is short for fold change).

For correlation analysis of gene expression between bulk RNA-seq datasets from protoplasted, un-protoplasted tissues and pseudo scRNA-seq datasets, the log2 (mean RPM+1) expression values were calculated for each gene and the Pearson-correlation coefficients were determined in R.

### Analysis and integration of scRNA-seq datasets

Raw data downloaded from NCBI were converted into FASTQ files by fastq-dump ^29^. After quality control, clean reads were mapped to reference genomes of plant species using CellRanger ^30^ with the default parameters. All downstream single-cell analyses were performed using Seurat (Stuart et al., 2019), which was implemented in R. In brief, the gene-cell matrices were loaded and then normalization was performed. In the principal component (PC) analysis, the scaled data were reduced to 50 PCs by setting “npcs = 50”. Then clusters were identified using the *FindClusters* function with the parameter of resolution set to 0.5. R package Harmony ^31^ were used to integrate datasets with the default parameter values. Finally, the *FindAllMarkers* function was used to identify marker genes that were up-regulated in each cluster versus all other cells (average logFC >= 0.25 plus maximum adjusted *p*-value ≤ 0.05). At the same time, the min.pct argument requires that genes are detected within a minimum percentage (i.e. 25%) of cells of a given cluster.

### GO enrichment analysis

Enriched GO terms associated with DEGs were identified by clusterProfiler version 4.0.5 ^32^. In detail, functions *compareCluster* and *gofilter* (level = 3) were used with the following parameters fun = “enrichGO”, OrgDb = “*db”, ont = “ALL”, pAdjustMethod = “BH”, pvalueCutoff = 0.05, qvalueCutoff = 0.05. The following GO annotation database packages were used: *A. thaliana* (OrgDb: org.At.tair.db), *Z. mays* (AnnotationHub: AH80549 | org.Zea_mays.eg.sqlite), *O. sativa* (OrgDb: org.OSativa. eg.db).

### BY-2 cell protoplast isolation

Protoplasts were prepared from 3 d-old tobacco BY-2 suspension cells. The 40-ml cell culture was filtered using a 40-μm nylon mesh (BD Falcon, USA) to collect BY2 cells, which were washed twice with protoplast wash solution (0.5 M mannitol, 4 mM MES (2-(N-morpholino) ethane sulfonic acid), pH 5.7). The cells were then divided into three parts. One part was immediately cryopreserved in liquid nitrogen for bulk RNA paired-end sequencing (with three replicates), the other two parts were resuspended in 10 mL of enzyme solution (pH 5.7), containing 1.5% cellulase “Onozuka RS” (Yakult Pharmaceutical Ind. Co. Ltd., Tokyo, Japan), 0.5% macerase (Yakult Pharmaceutical Ind. Co. Ltd., Tokyo, Japan), 0.5 M mannitol, and 4 mM MES, in a 20-ml Petri dish and incubated for 2 h at 26°C on a rotary shaker that is maintained in the dark at 60 rpm. Protoplasts were recovered by filtration through a 40-μm nylon mesh and washed twice with protoplast wash solution at 150 g (centrifugation) for 3 min. The obtained BY2 protoplasts were divided into two parts and cryopreserved, one for bulk RNA paired-end sequencing (with three replicates) and another for single-cell RNA sequencing (with two replicates). Illumina libraries were constructed with Gene Expression v3 kits (10X Genomics).

### Cell type annotation for scRNA-seq data from tobacco BY-2 cells

Cluster 4 were considered as cells in the S phase, exhibiting high expression of DNA-associated proteins and histones, such as genes from several HISTONE family members, including HIS4 (gene-65415, gene-65414), HIS3 (gene-44765), HIS2A (gene-24656), and HIS1 (gene-32296), which were essential for packaging nascent DNA into nucleosomes during DNA synthesis ^33^.

Cluster 5 corresponded to cells in the G2-M phase, marked by specific marker genes, such as CYCB1 (gene-36394, gene-6872) ^34^, MAD2 (gene-48930, gene-36942) ^35^, CSS52B (gene-75526), ENODL15 (gene-69508, gene-23349), and KNOLLE (gene-16029). GO enrichment analysis indicated that cluster 5 was enriched in pathways related to microtubule binding, mitotic cell cycle process, sister chromatid segregation, and nuclear division (**Table S4**).

Cluster 2 was classified as representing cells in the G0-G1 phase, characterized by high expression of PROLIFERATING CELL NUCLEAR ANTIGEN (PCNA, gene-18434) and THYMIDYLATE SYNTHASE (TS, gene-30912) genes associated with the G1/S transition ^36, 37^, marking the beginning of the cell cycle.

Cluster 3 showed characteristics of cell cycle exit, GO analysis supports the idea of cell cycle exit in cluster 3, with enrichment of terms associated with DNA damage and mitotic exit (**Table S4**). Genes such as TYROSYL-DNA PHOSPHODIESTERASE-RELATED (TDP1: gene-71104), MEDIATOR OF DNA DAMAGE CHECKPOINT PROTEIN 1’ (MDC1: gene-18581), and 5’-3’ EXORIBONUCLEASE 3 (XRN3: gene-4852) were specifically expressed in cluster 3 and linked to DNA damage response and cell cycle exit ^38, 39^.

Cluster 1 posed challenges for annotation due to the lack of specific cell cycle markers. The marker genes in cluster 1 exhibited features of both cell division and response to abiotic stress. Additionally, cluster 1 cells showed enrichment of pathways involved in abiotic stress responses, suggesting a mixture of cells arrested at different phase transitions in response to adverse environmental conditions (**Table S4**).

### Trajectory analysis

Pseudo-time trajectory analysis was performed using the algorithm of the Monocle2 version 2.20.0 ^40^ on the overall cluster marker gene identified by Seurat. Cells were ordered along the trajectory and visualized in a reduced dimensional space. Significantly changed genes along the pseudo-time were identified using the differential *GeneTest* function of Monocle2 with *q*-value < 0.01. Genes dynamically expressed along the pseudo-time were clustered using the *plotpseudotimeheatmap* function with the default parameters.

### Data and code availability

We collected publicly available datasets encompassing four types of omics data, including bulk RNA-seq, scRNA-seq, snRNA-seq, and spatial transcriptomics.

For the scRNA-seq datasets, the accession IDs are as follows: (1) *A. thaliana*: roots datasets PRJNA509920 ^17^, PRJNA640389 ^4^; leaves dataset ETAB-10016MT ^19^; PRJNA577177 ^41^; PRJNA678377 (Kim et al., 2021); floral receptacles PRJNA857332^42^. (2) *O. sativa*: roots dataset PRJNA706435 ^43^; roots and leaves dataset CRA004082^44^; inflorescence dataset ^45^; (3) *Z. mays*: leaves dataset ^23^; (4) *S. lycopersicum*: shoot borne roots ^8^.

For snRNA-seq datasets, the accession ID is PRJNA649267 from *A. thaliana* root^18^. The spatial transcriptomic datasets accession ID is CNP0002618 ^20^.

For bulk RNA-seq datasets, the accession IDs are *A. thaliana* root GSE123818 ^17^, leaf GSE161411 ^46^; *Z. mays* leaf GSE157758 ^23^, ear PRJNA647196 ^22^; *O. sativa* seedlings CRA004082 ^44^.

The scRNA-seq and bulk RNA-seq datasets generated from BY-2 cells in this study have been deposited in the NCBI Sequence Read Archive under the accession number GSE193131.

## Author contributions

**Jie Yao**: Writing - Original draft, Data curation, Methodology, Formal analysis, Visualization. **Nianmin Shang**: Resources, Software, Methodology. **Hongyu Chen**: Resources; Software. **Yurong Hu**: Resources; Software. **Qian-Hao Zhu**: Writing - Review & Editing, Supervision. **Longjiang Fan**: Writing - Review & Editing, Supervision. **Chuyu Ye**: Writing - Review & Editing, Supervision. **Qinjie Chu**: Writing - Review & Editing, Supervision, Investigation, Funding acquisition, Conceptualization, Project administration.

## Supplementary data

### Supplementary tables

**Table S1** List of plant bulk RNA-seq datasets used in this study before or after protoplasting

**Table S2** List of protoplasting induced differentially expressed genes (ppDEGs) in different plant species

**Table S3** Summary of publications involving plant scRNA-seq for considering protoplasting effects

**Table S4** GO enrichment results of tobacco ppDEGs or marker genes on each cluster on BY-2 scRNA-seq datasets

## Reference

1. Bawa, G., Liu, Z., Yu, X., Qin, A., and Sun, X. (2022). Single-Cell RNA Sequencing for Plant Research: Insights and Possible Benefits. International journal of molecular sciences 23. 10.3390/ijms23094497.

2. Liao, R.-Y., and Wang, J.-W. (2023). Analysis of meristems and plant regeneration at single-cell resolution. Current opinion in plant biology 74, 102378. 10.1016/j.pbi.2023.102378.

3. 3. Otero, S., Gildea, I., Roszak, P., Yipeng, L., Di Vittori, V., Bourdon, M., Kalmbach, L., Blob, B., Heo, J., and Peruzzo, F., et al. (2022). Research data supporting "A root phloem pole cell atlas reveals common transcriptional states in protophloem adjacent cells" (Apollo - University of Cambridge Repository).

4. Shahan, R., Hsu, C.-W., Nolan, T.M., Cole, B.J., Taylor, I.W., Greenstreet, L., Zhang, S., Afanassiev, A., Vlot, A.H.C., and Schiebinger, G., et al. (2022). A single-cell Arabidopsis root atlas reveals developmental trajectories in wild-type and cell identity mutants. Developmental cell 57, 543–560.e9. 10.1016/j.devcel.2022.01.008.

5. Han, E., Geng, Z., Qin, Y., Wang, Y., and Ma, S. (2024). Single-cell network analysis reveals gene expression programs for Arabidopsis root development and metabolism. Plant communications 5, 100978. 10.1016/j.xplc.2024.100978.

6. Yao, J., Chu, Q., Guo, X., Shao, W., Shang, N., Luo, K., Li, X., Chen, H., Cheng, Q., and Mo, F., et al. (2024). Spatiotemporal transcriptomic landscape of rice embryonic cells during seed germination. Developmental cell. 10.1016/j.devcel.2024.05.016.

7. 7. Wang, Q., Guo, Q., Shi, Q., Yang, H., Liu, M., Niu, Y., Quan, S., Di Xu, Chen, X., and Li, L., et al. (2024). Histological and single-nucleus transcriptome analyses reveal the specialized functions of ligular sclerenchyma cells and key regulators of leaf angle in maize. Molecular plant 17, 920–934. 10.1016/j.molp.2024.05.001.

8. Omary, M., Gil-Yarom, N., Yahav, C., Steiner, E., Hendelman, A., and Efroni, I. (2022). A conserved superlocus regulates above- and belowground root initiation. Science (New York, N.Y.) 375, eabf4368. 10.1126/science.abf4368.

9. Yue, H., Chen, G., Zhang, Z., Guo, Z., Zhang, Z., Zhang, S., Turlings, T.C.J., Zhou, X., Peng, J., and Gao, Y., et al. (2024). Single-cell transcriptome landscape elucidates the cellular and developmental responses to tomato chlorosis virus infection in tomato leaf. Plant, cell & environment 47, 2660–2674. 10.1111/pce.14906.

10. Zhang, Y., Chen, S., Xu, L., Chu, S., Yan, X., Lin, L., Wen, J., Zheng, B., Chen, S., and Li, Q. (2024). Transcription factor PagMYB31 positively regulates cambium activity and negatively regulates xylem development in poplar. The Plant cell 36, 1806–1828. 10.1093/plcell/koae040.

11. Bai, Y., Liu, H., Lyu, H., Su, L., Xiong, J., and Cheng, Z.-M.M. (2022). Development of a single-cell atlas for woodland strawberry (Fragaria vesca) leaves during early Botrytis cinerea infection using single cell RNA-seq. Horticulture research 9. 10.1093/hr/uhab055.

12. 12. Kang, M., Choi, Y., Kim, H., and Kim, S.-G. (2021). Single-cell RNA-sequencing of Nicotiana attenuata petal cells reveals the entire biosynthetic pathway of a floral scent.

13. Cervantes-Pérez, S.A., Thibivillliers, S., Tennant, S., and Libault, M. (2022). Review: Challenges and perspectives in applying single nuclei RNA-seq technology in plant biology. Plant science : an international journal of experimental plant biology 325, 111486. 10.1016/j.plantsci.2022.111486.

14. 14. Tian, C., Du, Q., Xu, M., Du, F., and Jiao, Y. (2020). Single-nucleus RNA-seq resolves spatiotemporal developmental trajectories in the tomato shoot apex. 10.1101/2020.09.20.305029.

15. Dolan, L., Janmaat, K., Willemsen, V., Linstead, P., Poethig, S., Roberts, K., and Scheres, B. (1993). Cellular organisation of the Arabidopsis thaliana root. Development (Cambridge, England) 119, 71–84. 10.1242/dev.119.1.71.

16. Hao, Y., Stuart, T., Kowalski, M.H., Choudhary, S., Hoffman, P., Hartman, A., Srivastava, A., Molla, G., Madad, S., and Fernandez-Granda, C., et al. (2024). Dictionary learning for integrative, multimodal and scalable single-cell analysis. Nature biotechnology 42, 293–304. 10.1038/s41587-023-01767-y.

17. Denyer, T., Ma, X., Klesen, S., Scacchi, E., Nieselt, K., and Timmermans, M.C.P. (2019). Spatiotemporal Developmental Trajectories in the Arabidopsis Root Revealed Using High-Throughput Single-Cell RNA Sequencing. Developmental cell 48, 840–852.e5. 10.1016/j.devcel.2019.02.022.

18. Farmer, A., Thibivilliers, S., Ryu, K.H., Schiefelbein, J., and Libault, M. (2021). Single-nucleus RNA and ATAC sequencing reveals the impact of chromatin accessibility on gene expression in Arabidopsis roots at the single-cell level. Molecular plant 14, 372–383. 10.1016/j.molp.2021.01.001.

19. 19. Tenorio Berrío, R., Verstaen, K., Vandamme, N., Pevernagie, J., Achon, I., van Duyse, J., van Isterdael, G., Saeys, Y., Veylder, L. de, and Inzé, D., et al. (2022). Single-cell transcriptomics sheds light on the identity and metabolism of developing leaf cells. Plant physiology 188, 898–918. 10.1093/plphys/kiab489.

20. Xia, K., Sun, H.-X., Li, J., Li, J., Zhao, Y., Chen, L., Qin, C., Chen, R., Chen, Z., and Liu, G., et al. (2022). The single-cell stereo-seq reveals region-specific cell subtypes and transcriptome profiling in Arabidopsis leaves. Developmental cell 57, 1299–1310.e4. 10.1016/j.devcel.2022.04.011.

21. Kumpf, R.P., and Nowack, M.K. (2015). The root cap: a short story of life and death. Journal of experimental botany 66, 5651–5662. 10.1093/jxb/erv295.

22. Xu, X., Crow, M., Rice, B.R., Li, F., Harris, B., Liu, L., Demesa-Arevalo, E., Lu, Z., Wang, L., and Fox, N., et al. (2021). Single-cell RNA sequencing of developing maize ears facilitates functional analysis and trait candidate gene discovery. Developmental cell 56, 557–568.e6. 10.1016/j.devcel.2020.12.015.

23. Bezrutczyk, M., Zöllner, N.R., Kruse, C.P.S., Hartwig, T., Lautwein, T., Köhrer, K., Frommer, W.B., and Kim, J.-Y. (2021). Evidence for phloem loading via the abaxial bundle sheath cells in maize leaves. The Plant cell 33, 531–547. 10.1093/plcell/koaa055.

24. Bakken, T.E., Hodge, R.D., Miller, J.A., Yao, Z., Nguyen, T.N., Aevermann, B., Barkan, E., Bertagnolli, D., Casper, T., and Dee, N., et al. (2018). Single-nucleus and single-cell transcriptomes compared in matched cortical cell types. PloS one 13, e0209648. 10.1371/journal.pone.0209648.

25. Williams, C.G., Lee, H.J., Asatsuma, T., Vento-Tormo, R., and Haque, A. (2022). An introduction to spatial transcriptomics for biomedical research. Genome medicine 14, 68. 10.1186/s13073-022-01075-1.

26. Dobin, A., Davis, C.A., Schlesinger, F., Drenkow, J., Zaleski, C., Jha, S., Batut, P., Chaisson, M., and Gingeras, T.R. (2013). STAR: ultrafast universal RNA-seq aligner. Bioinformatics (Oxford, England) 29, 15–21. 10.1093/bioinformatics/bts635.

27. Liao, Y., Smyth, G.K., and Shi, W. (2014). featureCounts: an efficient general purpose program for assigning sequence reads to genomic features. Bioinformatics (Oxford, England) 30, 923–930. 10.1093/bioinformatics/btt656.

28. Love, M.I., Huber, W., and Anders, S. (2014). Moderated estimation of fold change and dispersion for RNA-seq data with DESeq2. Genome biology 15, 550. 10.1186/S13059-014-0550-8.

29. Leinonen, R., Sugawara, H., and Shumway, M. (2011). The sequence read archive. Nucleic acids research 39, D19–21. 10.1093/nar/gkq1019.

30. Zheng, G.X.Y., Terry, J.M., Belgrader, P., Ryvkin, P., Bent, Z.W., Wilson, R., Ziraldo, S.B., Wheeler, T.D., McDermott, G.P., and Zhu, J., et al. (2017). Massively parallel digital transcriptional profiling of single cells. Nature communications 8, 14049. 10.1038/ncomms14049.

31. Korsunsky, I., Millard, N., Fan, J., Slowikowski, K., Zhang, F., Wei, K., Baglaenko, Y., Brenner, M., Loh, P.-R., and Raychaudhuri, S. (2019). Fast, sensitive and accurate integration of single-cell data with Harmony. Nature methods 16, 1289–1296. 10.1038/s41592-019-0619-0.

32. Yu, G., Wang, L.-G., Han, Y., and He, Q.-Y. (2012). clusterProfiler: an R package for comparing biological themes among gene clusters. Omics : a journal of integrative biology 16, 284–287. 10.1089/omi.2011.0118.

33. Chabouté, M.-E., Clément, B., and Philipps, G. (2002). S phase and meristem-specific expression of the tobacco RNR1b gene is mediated by an E2F element located in the 5’ leader sequence. The Journal of biological chemistry 277, 17845–17851. 10.1074/JBC.M200959200.

34. Combettes, B., Reichheld, J.P., Chabouté, M.E., Philipps, G., Shen, W.H., and Chaubet-Gigot, N. (1999). Study of phase-specific gene expression in synchronized tobacco cells. Methods in cell science : an official journal of the Society for In Vitro Biology 21, 109–121. 10.1023/A:1009880705257.

35. Musacchio, A., and Hardwick, K.G. (2002). The spindle checkpoint: structural insights into dynamic signalling. Nature reviews. Molecular cell biology 3, 731–741. 10.1038/NRM929.

36. Nelms, B., and Walbot, V. (2019). Defining the developmental program leading to meiosis in maize. Science (New York, N.Y.) 364, 52–56. 10.1126/SCIENCE.AAV6428.

37. Zhang, T.-Q., Chen, Y., Liu, Y., Lin, W.-H., and Wang, J.-W. (2021). Single-cell transcriptome atlas and chromatin accessibility landscape reveal differentiation trajectories in the rice root. Nature communications 12, 2053. 10.1038/s41467-021-22352-4.

38. Pawlik, T.M., and Keyomarsi, K. (2004). Role of cell cycle in mediating sensitivity to radiotherapy. International journal of radiation oncology, biology, physics 59, 928–942. 10.1016/j.ijrobp.2004.03.005.

39. Veylder, L. de, Beeckman, T., and Inzé, D. (2007). The ins and outs of the plant cell cycle. Nature reviews. Molecular cell biology 8, 655–665. 10.1038/NRM2227.

40. Qiu, X., Mao, Q., Tang, Y., Wang, L., Chawla, R., Pliner, H.A., and Trapnell, C. (2017). Reversed graph embedding resolves complex single-cell trajectories. Nature methods 14, 979–982. 10.1038/NMETH.4402.

41. Liu, Z., Zhou, Y., Guo, J., Li, J., Tian, Z., Zhu, Z., Wang, J., Wu, R., Zhang, B., and Hu, Y., et al. (2020). Global Dynamic Molecular Profiling of Stomatal Lineage Cell Development by Single-Cell RNA Sequencing. Molecular plant 13, 1178–1193. 10.1016/j.molp.2020.06.010.

42. Taylor, I.W., Patharkar, O.R., Mijar, M., Hsu, C.-W., Baer, J., Niederhuth, C.E., Ohler, U., Benfey, P.N., and Walker, J.C. (2022). Arabidopsis uses a molecular grounding mechanism and a biophysical circuit breaker to limit floral abscission signaling. 10.1101/2022.07.14.500021.

43. Zhang, T.-Q., Chen, Y., Liu, Y., Lin, W.-H., and Wang, J.-W. (2021). Single-cell transcriptome atlas and chromatin accessibility landscape reveal differentiation trajectories in the rice root. Nature communications 12, 2053. 10.1038/s41467-021-22352-4.

44. Wang, Y., Huan, Q., Li, K., and Qian, W. (2021). Single-cell transcriptome atlas of the leaf and root of rice seedlings. Journal of genetics and genomics 48, 881–898. 10.1016/j.jgg.2021.06.001.

45. Zong, J., Wang, L., Zhu, L., Bian, L., Zhang, B., Chen, X., Huang, G., Zhang, X., Fan, J., and Cao, L., et al. (2022). A rice single cell transcriptomic atlas defines the developmental trajectories of rice floret and inflorescence meristems. The New phytologist 234, 494–512. 10.1111/nph.18008.

46. Kim, J.-Y., Symeonidi, E., Pang, T.Y., Denyer, T., Weidauer, D., Bezrutczyk, M., Miras, M., Zöllner, N., Hartwig, T., and Wudick, M.M., et al. (2021). Distinct identities of leaf phloem cells revealed by single cell transcriptomics. The Plant cell 33, 511–530. 10.1093/plcell/koaa060.

