## Supplemental Figures for "Sub-clustering, marker identification and multi-omics integration influenced by protoplasting in plant scRNA-seq data analysis"

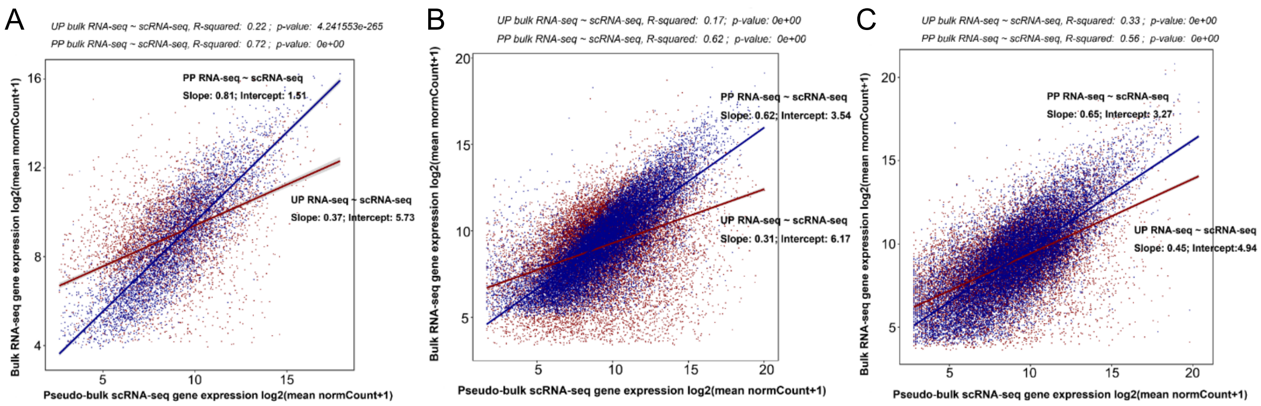


**Fig. S1** Correlation of gene expression values between scRNA-seq (pseudo bulk RNA-seq) and bulk RNA-seq data in *A. thaliana* **(A)**, *O. sativa* **(B)** and *Z. mays* **(C)**.

Blue and red lines represent un-protoplasted and protoplasted samples, respectively.

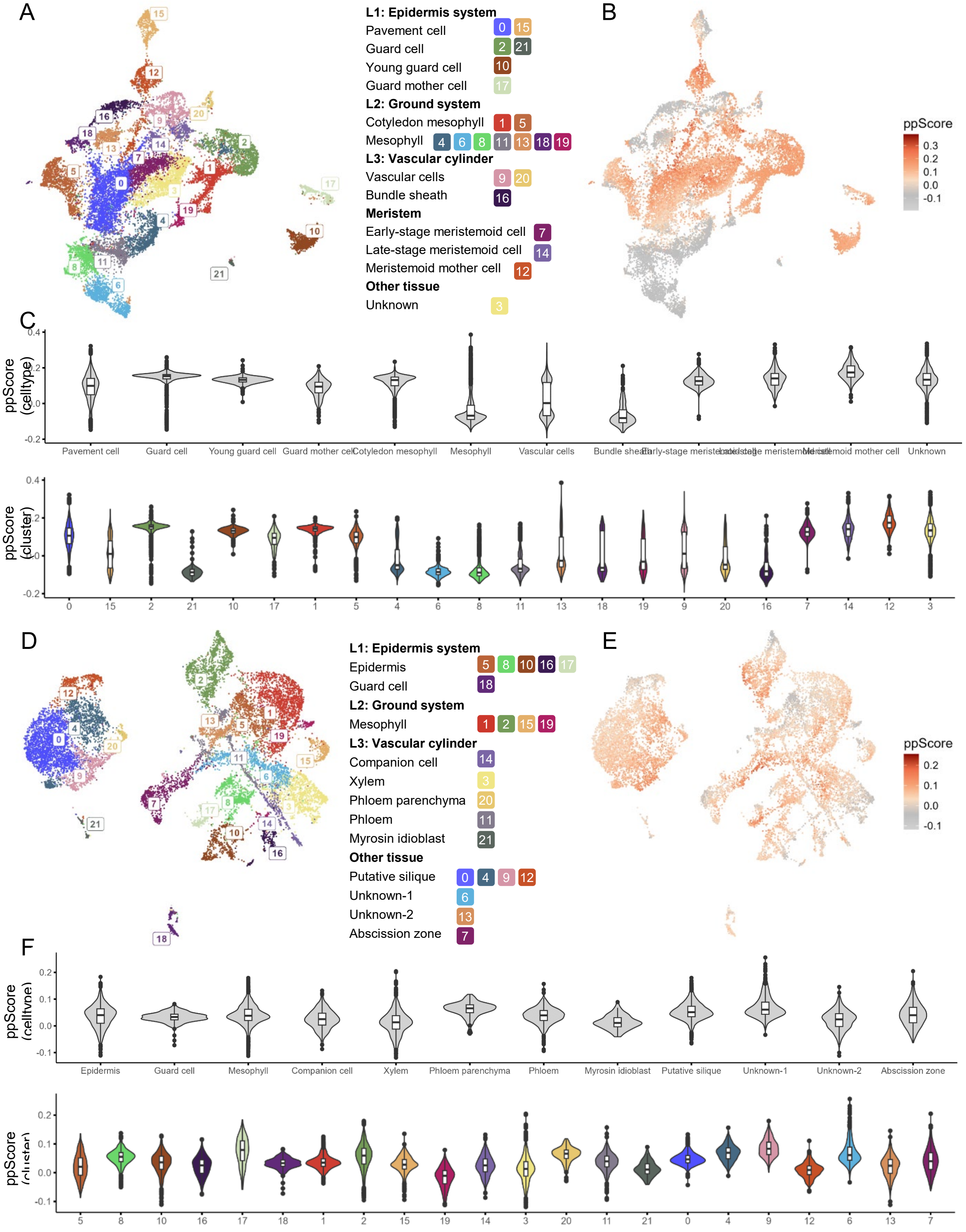


**Fig. S2** Protoplasting effects on *A. thaliana* leaf (A-C) and *A. thaliana* floral tissues (D-F).

**(A)** Clustering and annotated results of scRNA-seq data from *A. thaliana* leaf. (B) Display of ppScore for each single (C) Violin plots depicting ppScore values on cell types (upper) and clusters (down). The displayed figures D-F were analogous to the A-C.

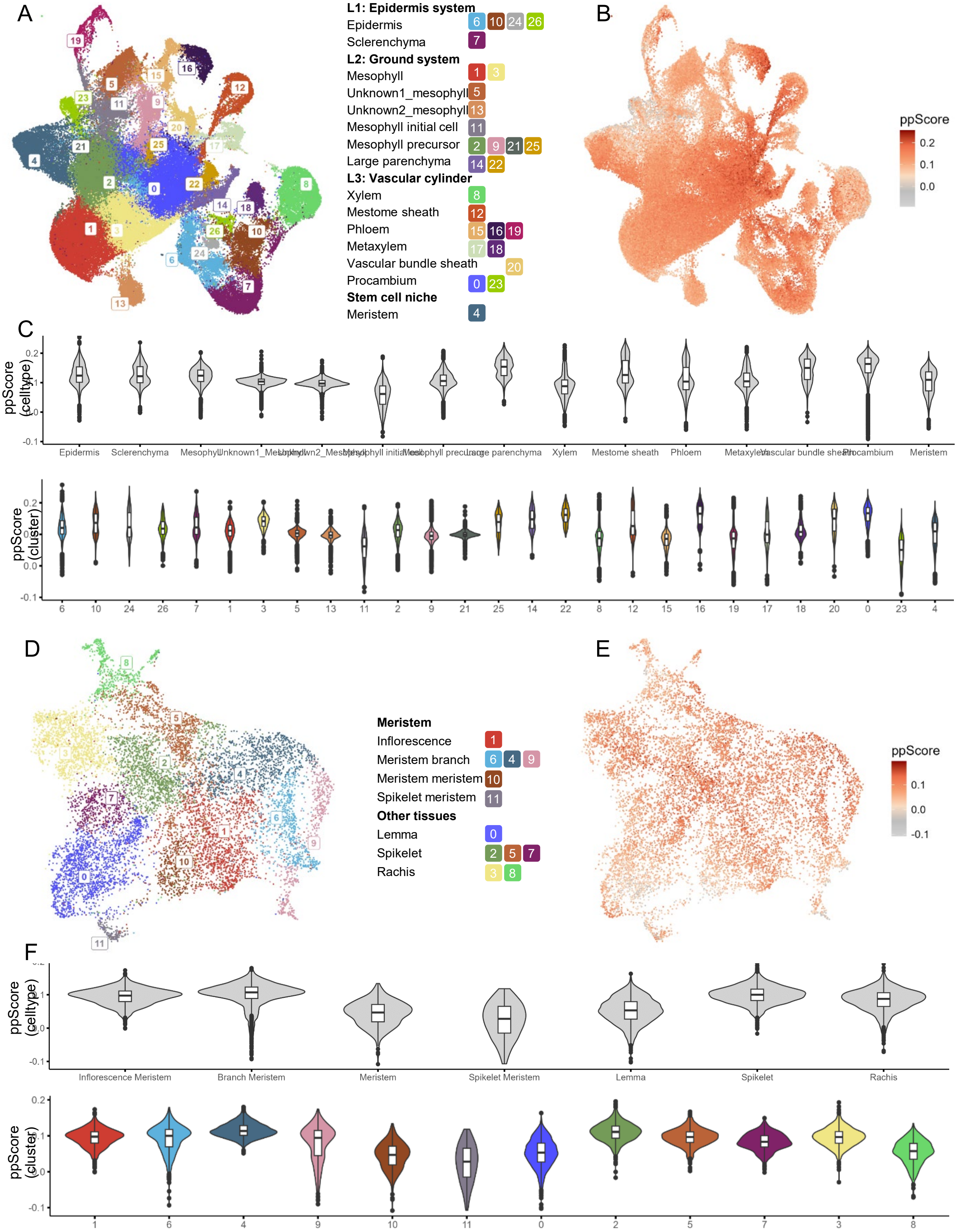


**Fig. S3** Protoplasting effects on *O. sativa* leaf (A-C) and *O. sativa* floral tissues (D-F).

**(A)** Clustering and annotated results of scRNA-seq data from *O. sativa* leaf. (B) Display of ppScore values for each single cell. **(C)** Violin plots depicting ppScore values on cell types (upper) and clusters (down). The displayed figures D-F were analogous to the A-C.

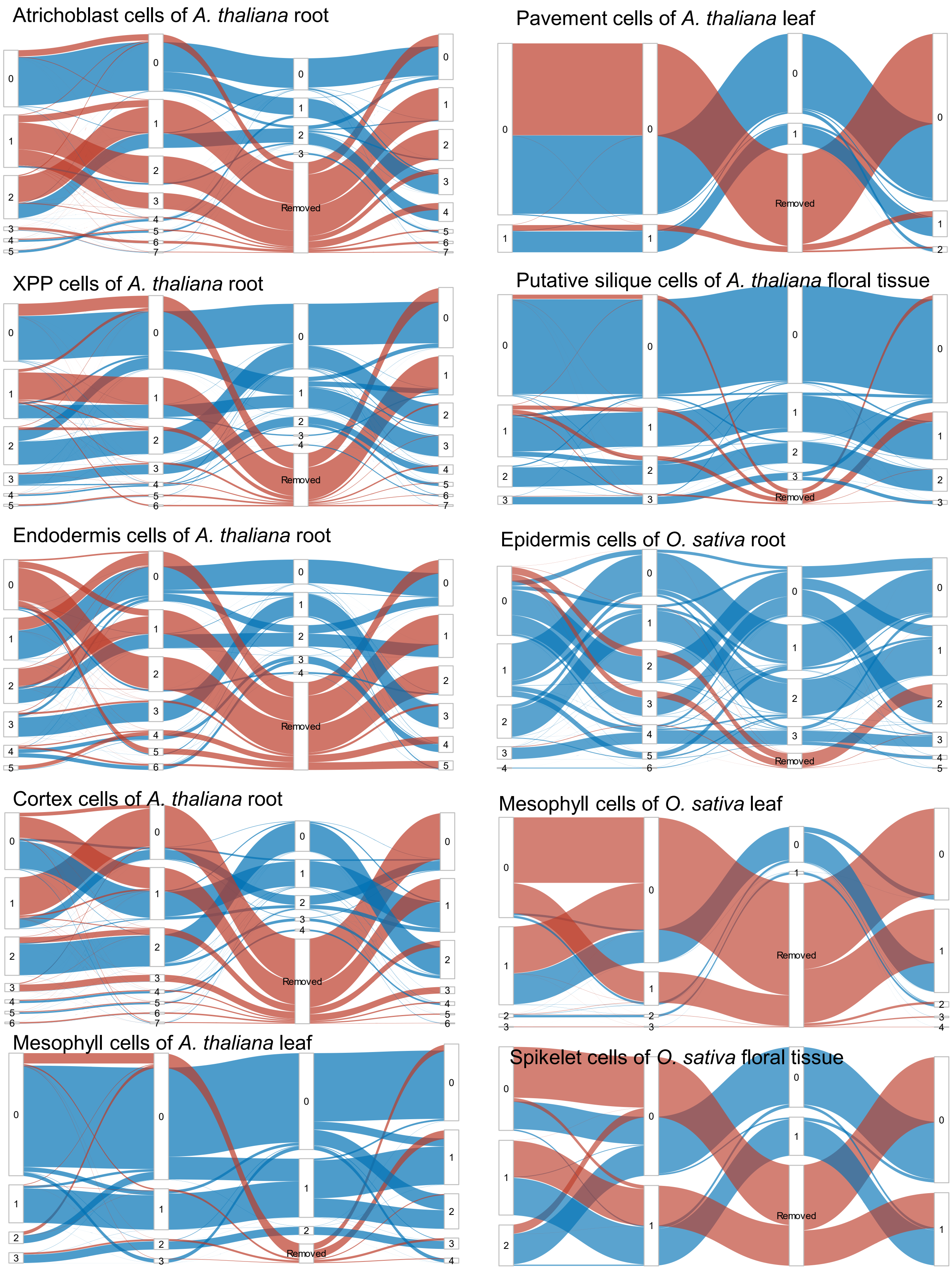


**Fig. S4** Clustering differences of different cell types between raw and modified datasets (i.e. removing protoplasting effects by three methods) with same parameters (res = 0.1).

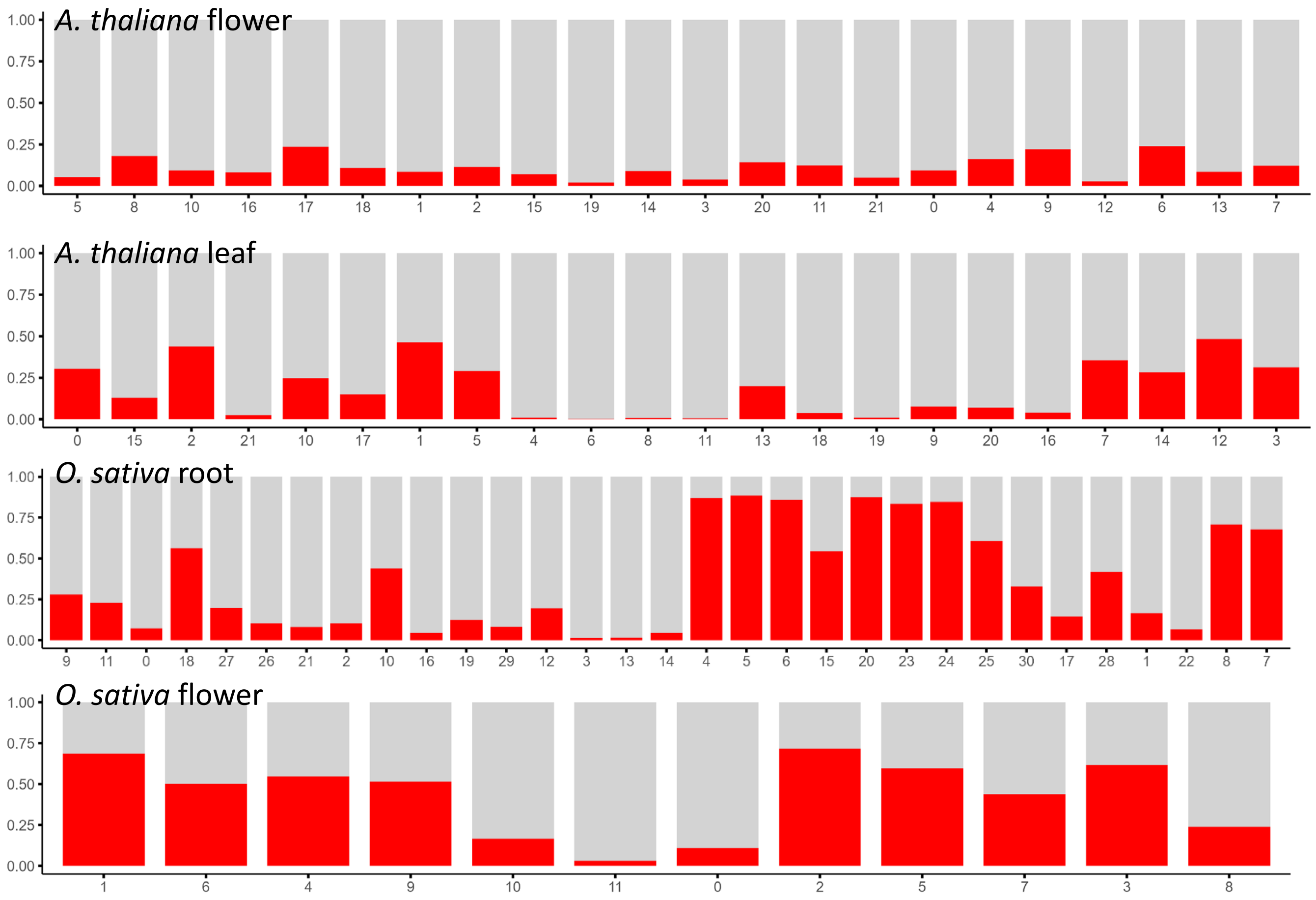


**Fig. S5** Proportion of protoplasting-related genes in marker genes of each cell cluster of different tissues. Red bars represent the proportion of ppDEGs while gray bars represent the other genes.

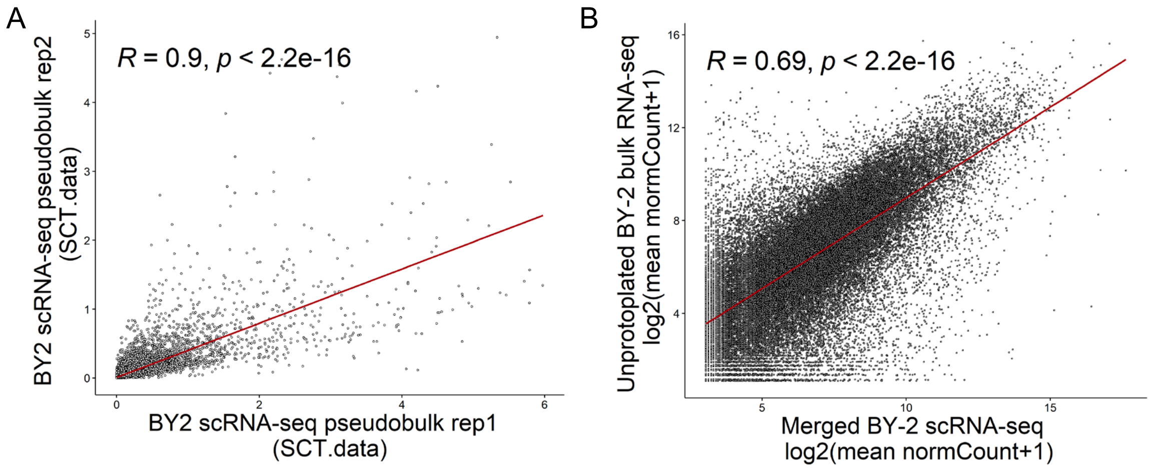


**Fig. S6** Characteristics of datasets from BY-2 cell lines generated in this study.

**A:** Comparison of similarity between two tobacco BY-2 scRNA-seq batches (Spearman's correlation = 0.90). **B:** Comparison of expression similarity between BY-2 scRNA-seq samples (protoplasted cells) and non-protoplasted samples (Spearman's correlation = 0.69), with each point representing a gene.
